# A semantic segmentation model to predict subcellular glycogen localization using transmission electron microscopy images

**DOI:** 10.64898/2026.02.04.703703

**Authors:** Anders A. Hansen, Jacob M. Egebjerg, Kristian Solem, Kristoffer J. Kolnes, Daniel Wüstner, Jørgen F. P. Wojtaszewski, Jørgen Jensen, Joachim Nielsen

**Author notes:** Corresponding author: Joachim Nielsen.

## Abstract

Transmission electron microscopy (TEM) is the gold standard for assessing subcellular glycogen localization in skeletal muscle fibres, but conventional manual analysis is extremely time-consuming and limits large-scale studies. Here, we developed and validated a deep learning–based semantic segmentation approach to automate quantification of glycogen particles across defined subcellular compartments in human skeletal muscle. Skeletal muscle biopsies were obtained from seven healthy men under conditions of normal, depleted, and supercompensated glycogen content. TEM images were acquired from myofibrillar and subsarcolemmal regions and manually annotated to train two complementary attention U-Net models: a region model identifying subcellular structures (intermyofibrillar space, intramyofibrillar regions including A-band, I-band and Z-disc, and mitochondria) and a glycogen model detecting individual glycogen particles. Combining the two models enabled estimation of compartment-specific glycogen areal densities. Model performance was evaluated against manual point-counting. At the fibre level, estimates based on 10–12 images per region achieved biases below 15% and coefficient of variation below 26% for all compartments. Importantly, model-derived total glycogen volume density showed strong concordance with biochemically determined muscle glycogen content across biopsies. In conclusion, this validated semantic segmentation workflow provides a robust, objective, and highly time-efficient tool for quantifying subcellular glycogen distribution in skeletal muscle. The model substantially reduces analysis time and enables high-throughput investigations of compartmentalized glycogen metabolism, with model weights and code made openly available.

## Introduction

Glycogen particles are the primary storage form of glucose in skeletal muscle fibres (Ball et al. 2011). They consist of branched chains of glucose units (glycosyl units) and typically measure between 20 and 30 nm in diameter, corresponding to approximately 5,000–10,000 glycosyl units per particle (Shearer & Graham, 2004). Their distribution within the fibre is uneven, with the majority (approximately 80%) located in the intermyofibrillar compartment, adjacent to the sarcoplasmic reticulum and mitochondria (Marchand et al. 2002). The remaining particles are distributed equally between the intramyofibrillar compartment, embedded within the myofibrils, and the subsarcolemmal compartment, situated near the fibre surface.

The subcellular distribution of glycogen can be assessed using transmission electron microscopy, applying refined imaging techniques to enhance glycogen contrast (De Bruijn, 1974; Marchand et al. 2002).

There is vast evidence for compartmentalized glycogen turnover in skeletal muscle (i.e. Marchand et al. 2007; Nielsen et al. 2022; Jensen et al. 2020; Schytz et al. 2024) and therefor an unmet need to perform studies on the role and regulation of subcellular glycogen localization. Research is limited, however, by the extreme time consumption for image analyses accumulating to around 1-2 hours per muscle fiber or 10-15 hours per biopsy (based on 8-10 fibres).

Inspired by recent advances in bioimaging, where automated segmentation has been successfully applied to predict mitochondria and lipid droplets (Bleck et al. 2018; Conrad & Narayan, 2023), we aimed to develop an AI model capable of identifying glycogen within subcellular regions of skeletal muscle fibres to save hundreds of working hours for image analyses in a typical project. During the development process, a new AI model demonstrating high accuracy was published (Ríos et al. 2024). Our model closely resembles this recently introduced approach. Notably, our model provides several important additional achievements beyond the prediction of intermyofibrillar and intramyofibrillar glycogen, including predictions of subsarcolemmal glycogen, Z-discs, and mitochondria. Furthermore, our model includes validation against biochemically determined glycogen concentrations, which was lacking in the model developed by Ríos et al.

## Methods

### Ethical approval

This study was approved by the Ethics Committee at the Norwegian School of Sport Sciences (ref. no. 40-191217) and reported to the Norwegian Centre for Research Data (ref. no. 858276). Oral and written informed consent were obtained from all participants before the start of the investigation.The study was conducted in accordance with the Declaration of Helsinki, except for pre-trial registration in a publicly available database

### Participants and muscle samples

From a larger on-going project, we included 21 skeletal muscle biopsies with varying glycogen content (to develop a model which would be robust over a wide range of glycogen levels). These biopsies originated from seven moderately trained men (means ± SEM; age: 30 ± 3 yr; weight: 73 ± 3 kg; height: 181 ± 2 cm), who had performed intermittent one-legged cycling until exhaustion followed by three days of recovery with a high carbohydrate intake. The skeletal muscle biopsies were obtained from vastus lateralis of the non-exercising leg before the exercise (normal glycogen), from the exercising leg both immediately after the exercise (low glycogen) and after the three-days recovery (high glycogen). Samples were processed for transmission electron microscopy and frozen for subsequent biochemical analyses.

### Biochemical determination of glycogen

Muscle samples were freeze-dried for at least 48 hours before being carefully cleaned of any visible fat, blood, and connective tissue, and subsequently homogenized. Detailed procedures for tissue preparation are described in Kolnes et al. (2025). Glycogen concentrations were assessed twice using distinct analytical approaches. The initial measurement employed a Pentra C400 analyzer (Horiba Medical, Triolab, Denmark) on 60 µl of homogenate, following acid hydrolysis by boiling in 2 mM HCl for two hours. To validate the unexpectedly low glycogen values obtained, a second analysis was performed using a fluorometric method with a Fluoroskan instrument (Thermo Fisher Scientific, Waltham, MA, USA), based on the protocol by Lowry & Passonneau (1972).

### Transmission electron microscopy

Muscle biopsy samples were initially fixed in 2.5% glutaraldehyde buffered with 0.1 M sodium cacodylate (pH 7.3) at 5°C. The preparation for transmission electron microscopy (TEM) followed procedures previously described (see Jensen et al. 2022). TEM was used to visualize and quantify glycogen distribution within individual muscle fibres. After a 24-hour fixation period, samples were rinsed multiple times in the same buffer and stored at 5°C until further processing. Post-fixation was carried out using a solution containing 1% osmium tetroxide and 1.5% potassium ferrocyanide in 0.1 M sodium cacodylate buffer for 90 minutes at 4°C. The tissue was then rinsed, dehydrated through a graded ethanol series (4–20°C), and infiltrated with increasing concentrations of propylene oxide and Epon resin at 20°C. Final embedding was done in pure Epon at 30°C.

Ultrathin longitudinal sections (60 nm) were cut using a Leica Ultracut UCT ultramicrotome and stained with uranyl acetate and lead citrate to enhance contrast. The sections were imaged using a calibrated Philips CM100 transmission electron microscope equipped with an Olympus Veleta digital camera at 66,000× magnification (0.77 nm/pixel).

For the manual segmentation and training of the model, images were obtained from sections originating from 12 different biopsies (4 participants) obtained from the control, exercise, or recovery leg. The distribution between legs ensured images with vast differences in glycogen content with the purpose to train a robust model across different glycogen levels. Images were acquired in a randomized, systematic manner: 24 from the subsarcolemmal region and 12-13 each from the superficial and central areas of the myofibrillar region.

For investigation of how the model’s prediction of total glycogen volume fraction associates with biochemical determination of glycogen, images from all 21 biopsies were included comprising 7-9 fibres per biopsy.

### Image analysis by semantic segmentation

The goal of the model was to predict the areal distribution of glycogen particles across three distinct subcellular compartments. This was achieved by developing a semantic segmentation model capable of identifying pixels corresponding to specific subcellular regions (region model) and glycogen particles (glycogen model). The approach closely resembles the model described by Ríos et al. (2024). By combining the outputs of both models (region and glycogen models), glycogen areal percentages were estimated within each subcellular compartment.

#### Region model

The region model was trained on 859 images exclusively from the myofibrillar compartment, excluding those from the subsarcolemmal region. These images were evenly distributed across pre-exercise, post-exercise, and three-day recovery biopsies from five participants, ensuring a representative dataset with varying glycogen content. Each image was partially annotated using the platform slicer.org (Fedorov et al. 2012), with only regions that could be confidently identified included in the annotation. Five predefined classes were used: A-band, I-band, Z-disc, intermyofibrillar space, and mitochondria. The A-band, I-band, and Z-disc collectively represent the intramyofibrillar compartment.

The region model utilized an attention U-Net architecture (Oktay et al. 2018) with an encoder-decoder structure employing progressively increasing filter sizes of [8, 16, 32, 64, 128, 256, 512, 1024] through the contracting path, with corresponding decreasing filter sizes in the expanding path. At the final decoder layer with 8 filters, the architecture incorporated dual output heads: a segmentation head employing softmax activation for multi-class prediction and a reconstruction head for auxiliary feature learning supervision.

The composite loss function combined multiple objectives to optimize both segmentation accuracy and feature representation: Dice loss and cross-entropy focal loss for segmentation, supplemented by mean squared error (MSE) loss for the reconstruction component weighted at 0.1. Training was conducted using Automatic Mixed Precision (AMP) with 16-bit precision to optimize memory usage and computational efficiency, employing the Adam optimizer for gradient-based parameter updates. The training dataset was augmented using geometric transformations including rotations, translations, scaling, and elastic deformations to enhance model generalization and robustness across varying image characteristics.

The performance of the trained region model is illustrated in Fig. 1, using examples from images with both high and low glycogen areal percentages. Although the model was trained exclusively on myofibrillar images, it was also applied to subsarcolemmal images. By visual inspection of several images, it was clear that the model mainly predicts subsarcolemmal glycogen as the intermyofibrillar class. Therefore, using the subsarcolemmal images, the intermyofibrillar class was relabelled as subsarcolemmal, and the A-band, I-band, and Z-disc classes were merged into a single category referred to as the non-subsarcolemmal (Non-SS) region. Subsarcolemmal images were captured in a manner that minimized inclusion of the intermyofibrillar region and avoided large portions of adjacent fibres. Therefore, it is assumed that the predicted intermyofibrillar region in these images corresponds to the subsarcolemmal compartment. Figure 2 presents three representative examples: a typical subsarcolemmal image, an image containing a nucleus, and a series of aligned images capturing an extended subsarcolemmal region. Model accuracy was evaluated using 92 images that were not part of the training dataset. These images were fully annotated by an experienced investigator (J.N.).

**Fig. 1.**
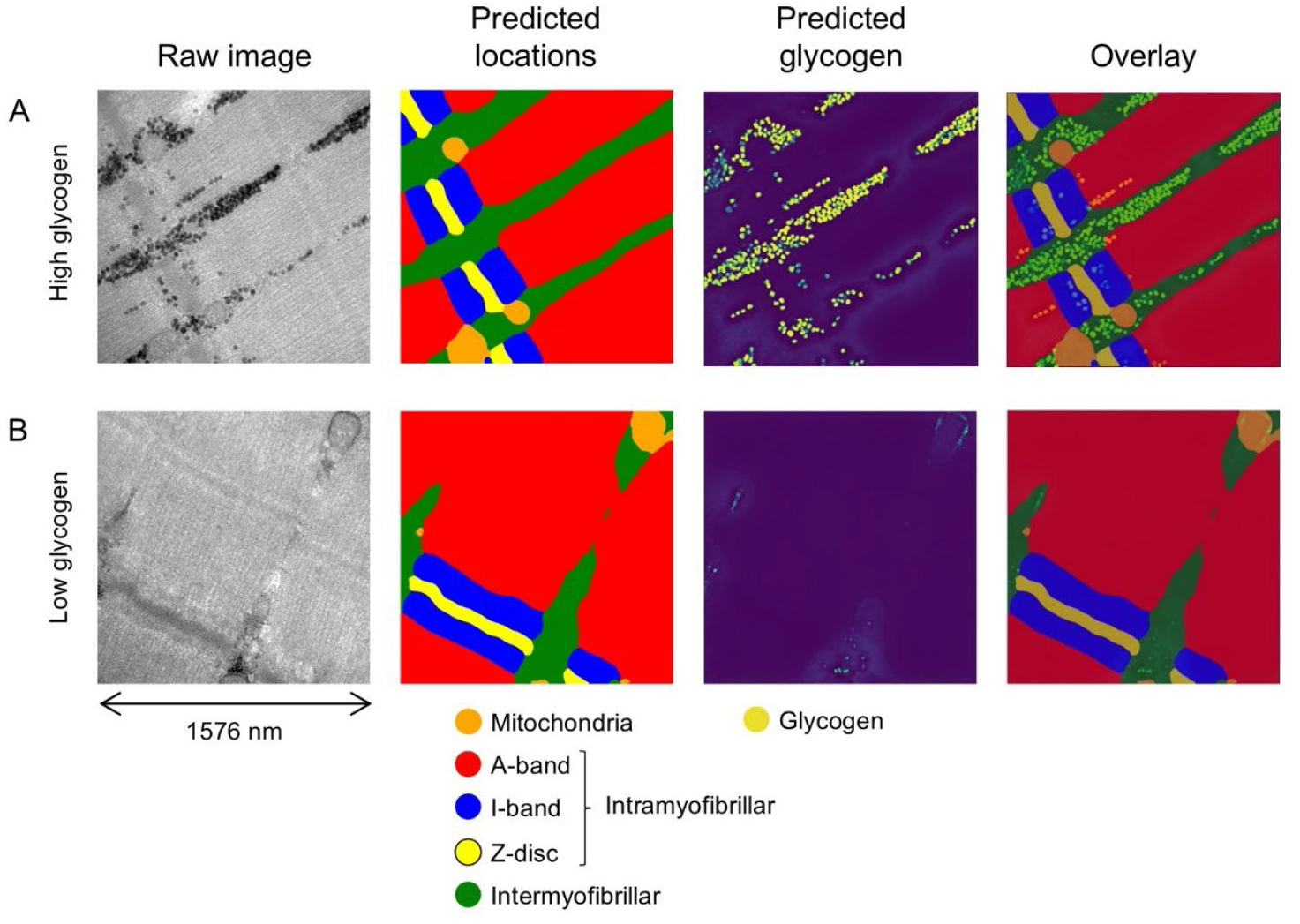
Illustration of the model used to predict glycogen in myofibrillar regions. A) Representative image with high glycogen content from a pre-exercise biopsy. B) Representative image with low glycogen content from a post-exercise biopsy.

**Fig. 2.**
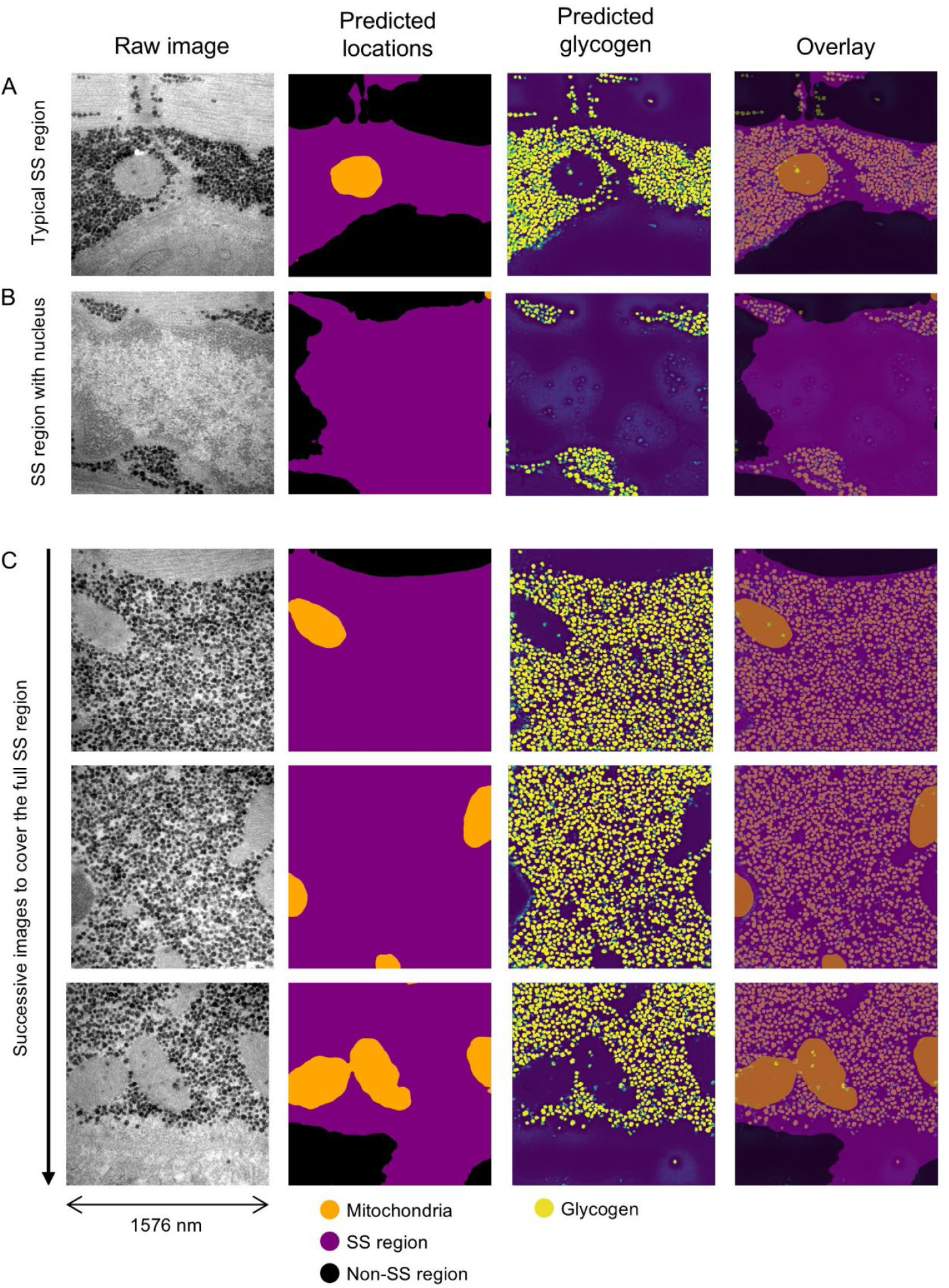
Illustration of the model used to predict glycogen in the subsarcolemmal region. A) Representative image with high glycogen content from a pre-exercise biopsy. B) Image containing a nucleus, demonstrating that the electron-dense nuclear material is not misclassified as glycogen. C) A series of aligned images captured to encompass the entire subsarcolemmal region. In some fibres, this region can extend several micrometres in depth (occasionally up to ~5 µm) requiring multiple images to fully cover the area.

#### Glycogen model

The glycogen model was constructed using a structure nearly identical to that of the region model, but with a binary classification distinguishing between glycogen and non-glycogen pixels. Training was performed on a subset of 100 images from the region model dataset, with equal representation from pre-exercise, post-exercise, and recovery time points to ensure inclusion of both small and large glycogen particles. The training process incorporated the same augmentation and variation techniques used for the region model. The model was evaluated using a probability threshold of 0.3 and the same 92 test images used for evaluating the region model. The manual estimation of glycogen areal fraction was performed by point-counting using a 37.4 × 37.4 nm grid (Jensen et al. 2022).

#### The combined model (glycogen areal percentages per region)

Validation of the combined model was performed using the same 92 myofibrillar images and an additional 84 randomly selected subsarcolemmal images, evenly distributed across all time points. Ground truth estimates were obtained through manual point counting, using grid sizes of 37.4 nm, 18.7 nm, and 74.8 nm for the intermyofibrillar, intramyofibrillar, and subsarcolemmal compartments, respectively, adjusted to match particle density.

#### Glycogen particle size, glycogen volume percentage, and numerical density

Isolated glycogen particles were extracted from the glycogen model predictions by applying a circularity threshold above 0.8 and selecting particles within a size range of 10 to 42 nm. These particles were used to estimate glycogen particle size and to calculate glycogen volume percentage. The volume percentage was derived by adjusting the areal percentage for particle size, assuming spherical geometry (Melendez et al. 1998), and accounting for the section thickness (60 nm).Calculations were based on the formula described by Weibel (1980): *V*V = *A*A - *t* [(1/*π*)·*B*A – *N*A·[(*t*·*H*)/(*t*+*H*)]], where *A*A is the glycogen area fraction, *t* is the section thickness, *B*A is the glycogen boundary length density, *N*A is the number of particles per area, and *H* is the mean glycogen particle diameter. Intramyofibrillar glycogen was expressed as volume per intramyofibrillar space, intermyofibrillar glycogen as per volume per myofibrillar space (inter- and intramyofibrillar), and subsarcolemmal glycogen as per surface area of the fibre as estimated by the image (1,6 µm) width multiplied by the section thickness (60 nm).

To estimate the total glycogen volume density across compartments including the subsarcolemmal compartment (expressed per surface area), muscle fibres were modelled as cylindrical structures with a diameter of 80 μm. Since glycogen in the intermyofibrillar and intramyofibrillar compartments was expressed as volume density, but subsarcolemmal glycogen as volume per unit surface area, a conversion of subsarcolemmal glycogen was necessary to enable comparison. Subsarcolemmal glycogen was recalculated to volume density (per fiber with a diameter of 80 µm) using the formula for the volume beneath the surface of a cylinder: (*V*b)=*R*×0.5×*A*, where *R* is fibre radius and *A* is the fibre surface area. Then, total glycogen volume density of each fiber was given by the formula: Total glycogen volume density = Intermyofibrillar glycogen volume density + intramyofibrillar glycogen volume density x intramyofibrillar volume density + subsarcolemmal glycogen / *V*b. Images from the superficial region of the myofibrillar space were weighted three times more than images from the central part of the myofibrillar space to take into account the higher contribution of the superficial region to the total.

### Data handling and statistics

Assessment of the model’s performance was conducted by analysing a confusion plot based on the 92 test images. Agreement between the model’s predictions and the ground truth (J.N.’s predictions) was assessed by Bland-Altman plots (Bland & Altman, 1986). For correlation and concordance analyses, statistical significance was defined as α ≤ 0.05.

## Results

### Image Analysis by Semantic Segmentation

#### Region model

Representative images and model predictions are shown in Fig. 1 and 2. Validation was performed using 92 images not included in the training dataset.

The model’s accuracy in predicting the two major regions of intermyofibrillar and intramyofibrillar was 91% and 93%, respectively. While the positive predictive value (PPV) was high for the intramyofibrillar region, it was only 81% for the intermyofibrillar region, where 16% of the predictions should have been intramyofibrillar (Fig. 3A). This misclassification led to an underestimation of the intramyofibrillar region and since this was skewed towards the A-band with very little glycogen, the glycogen areal percentage is overestimated most likely due to an underestimation of the reference space (intramyofibrillar region).

**Fig. 3.**
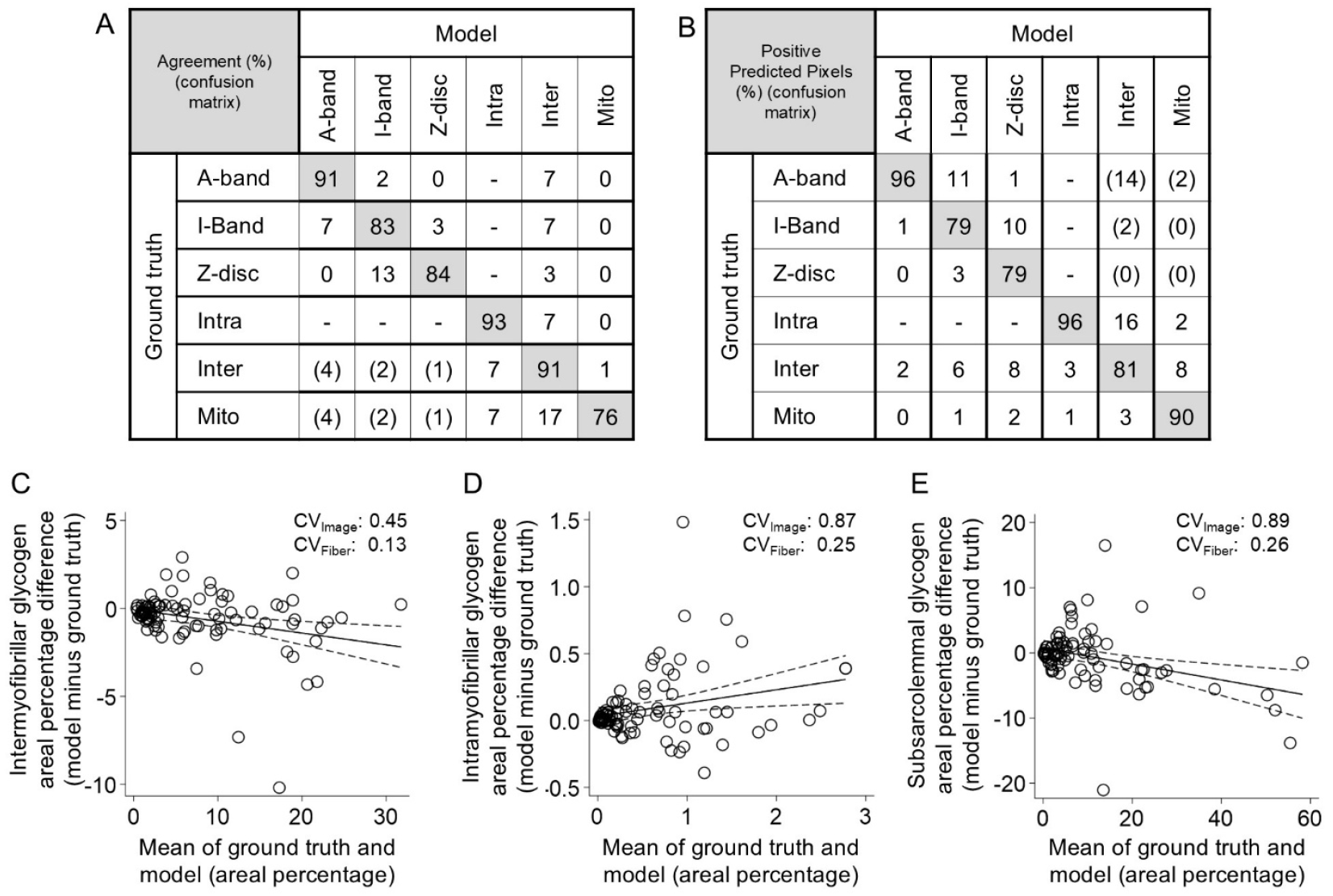
Performance of the model. A) Agreement between model predictions and ground truth is shown as the percentage of true positive pixels (grey boxes) relative to the total number of ground truth positive pixels. False negatives are shown in white boxes as the percentage of ground truth positive pixels not identified by the model. Numbers in parentheses represent a breakdown of the intramyofibrillar region into its subregions. B) Positive predictive values are shown as the percentage of true positive pixels relative to the total number of predicted positive pixels (grey boxes). False positives are shown in white boxes as the percentage of predicted positive pixels that were not present in the ground truth. Numbers in parentheses represent a breakdown of the intramyofibrillar region into its subregions. C-E). Bland–Altman plots showing the difference between ground truth and model-predicted glycogen areal percentages, plotted against the mean of the two values for each region. Linear regression revealed systematic biases: (intermyofibrillar glycogen) y = -0.067x-0.063 (R^2^=0.09; *P* = 0.005; N = 92); (intramyofibrillar glycogen) y = 0.0996x-0.0311 (R^2^=0.07; *P* = 0.012; N = 92); (subsarcolemmal glycogen) y = -0.122x-0.718 (R^2^=0.12; *P* = 0.002; N = 84).

Lower accuracy was observed for the smaller regions, including the I-band, Z-disc, and mitochondria (Fig. 3A). The confusion matrix revealed that Z-discs were frequently misclassified as I-band, I-band as A-band or intermyofibrillar, and mitochondria primarily as intermyofibrillar. Positive predictive values (PPVs) were high for A-band, intramyofibrillar, and mitochondria, but lower for the intermyofibrillar region due to overlap with intramyofibrillar predictions. PPVs for Z-disc and I-band were also reduced, with frequent misclassification as I-band and A-band, respectively.

#### Glycogen model

Model performance is illustrated in Fig. 1 and Fig. 2. When applied to the same 92 test images used for evaluating the region model, the glycogen model showed a 6% underestimation in areal percentage. The coefficient of variation (CV) across individual images was 0.42, which corresponds to a CV of 0.12 at the fibre level when averaging across 12 images per region.

#### Combined model

Across all regions, discrepancies between model predictions and ground truth were dependent on the areal percentage of glycogen. In general, the model tended to underestimate glycogen content in the intermyofibrillar and subsarcolemmal regions by 5–10%, while overestimating intramyofibrillar glycogen by 10–15% (Fig. 3).

To estimate glycogen localization at the single-fibre level, we examined the relationship between the number of images analysed and the uncertainty of the estimate in a subset of fibres sampled across all experimental time points. The data revealed a power-relationship: estimates based on only two images differed by 20–50% from those based on 24 images (Fig. 4A–C). This difference decreased substantially with more images, reaching 5–15% at 10–12 images.

**Fig. 4.**
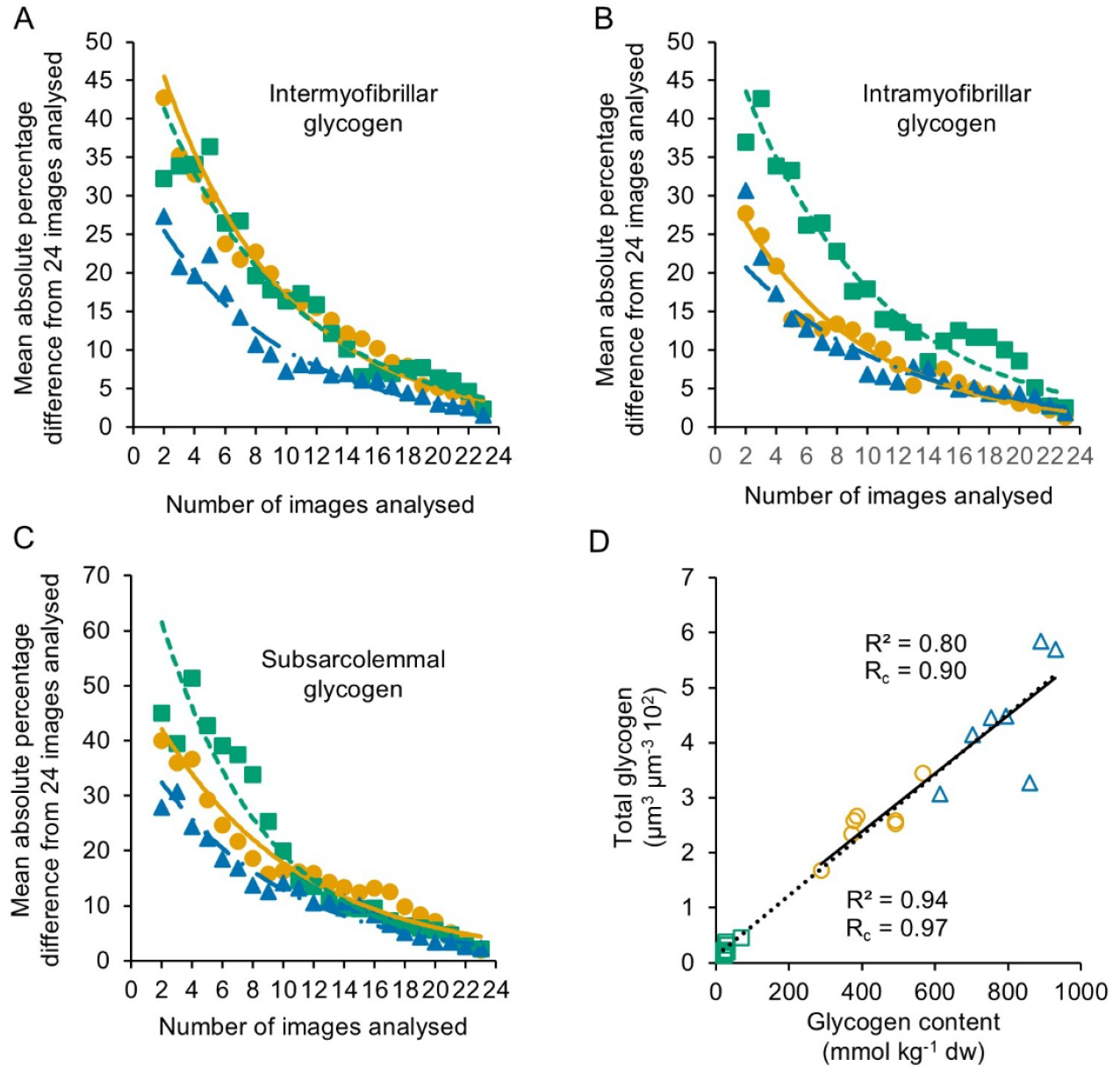
Relationship between number of images analysed and precision of the estimate. A-C) Plots show the mean absolute difference (%) between estimates based on 2–23 images and those based on 24 images, across 16, 10, and 14 fibres from pre-exercise (orange), post-exercise (green), and recovery (blue) biopsies, respectively. After analysing 12 images, the estimates differed from the 24-image reference by approximately 10–15% for intermyofibrillar and intramyofibrillar glycogen areal percentages, and by 15–20% for subsarcolemmal glycogen. D) The models prediction of total glycogen volume density showed high correlation (R^2^) and concordance (R_c_) with biochemical determined glycogen content. This is shown across all samples (dotted line, N = 21) and with post-exercise samples excluded (solid line, N = 14).

The average total glycogen volumetric content across 8–9 fibres per biopsy showed a strong correlation and concordance with chemically measured glycogen content from the same biopsy (Fig. 4D).

## Discussion

Here we developed a semantic segmentation model to automate the analysis of subcellular glycogen localization in skeletal muscle fibers based on transmission electron microscopy (TEM) images. The model demonstrated >90% agreement with manual image assessments.

The model developed in this study was built using similar methods and with comparable performance levels as the model by Rios et al. (2024). While the model from Rios et al. was unable to distinguish between Z-discs and mitochondria, our model successfully discriminated between these structures, enabling the prediction of both parameters and, in turn, allowing investigation of their associations with glycogen-related metrics. This capability may have important implications, as glycogen metabolism during exercise and recovery is strongly influenced by motor unit properties and muscle fibre types (Gollnick, 1974; Tsintzas et al. 1995), which are known to correlate with Z-disc width and mitochondrial content (Sjöström et al., 1982a, 1982b). Another key difference from the model by Rios et al. lies in the assessment of glycogen particle size. While Rios et al. simulated the effect of particle size on the probability of being intersected by the sectioning plane, we extracted all isolated particles (identified based on sphericity) and measured their area directly. We therefore assume that the extracted particles do not differ in size from those clustering closely together. This assumption is supported by the observation that the mean particle size in the present study aligns well with our previous findings using a manual method that included both isolated and clustered particles (Jensen et al., 2021; Hokken et al., 2021).

The model underestimated glycogen content in the intermyofibrillar and subsarcolemmal regions, while it overestimated content in the intramyofibrillar region. The underlying cause of this bias remains unclear. A common feature of the intermyofibrillar and subsarcolemmal regions is the clustering of glycogen particles, which may suggest that the model has limited ability to fully detect the area occupied by particle clusters. The overestimation of intramyofibrillar glycogen may be due to an underestimation of the A-band, as indicated by the confusion matrix (Fig. 3B). Since the A-band is relatively glycogen-poor, a reduced reference space combined with unchanged glycogen content could result in a slight overestimation of the volume fraction of glycogen in the intramyofibrillar region. We recommend that application of the model include a manual-to-model variation test, with any discrepancies corrected accordingly. Additionally, our experience indicates that the contrast of glycogen and mitochondria may vary between sample batches, further underscoring the need for careful and context-specific use of the model.

Taken together, we believe that our model represents a significant advancement in reducing the workload associated with image analysis. Our previous manual workflow required approximately 1.5 hours of imaging and 12 hours of analysis per biopsy. With the current model, this can be reduced to 2.5 hours of imaging and just 0.5 hours of analysis per biopsy saving roughly 220 hours of work in a project of similar scale (21 biopsies × 10.5 hours saved). In larger studies involving 60–100 biopsies (Jensen et al., 2020; Schytz et al., 2024), the reduction in workload becomes massive. Beyond its practical utility, the model also highlights the potential of deep learning in bioimaging. Further refinement of the method could uncover novel aspects of compartmentalized glycogen metabolism, such as particle clustering characteristics, single-particle alignment, and organelle–glycogen interactions, features that are not currently captured by the model.

In conclusion, the semantic segmentation model developed in this study provides a valid approach for predicting glycogen particles in subcellular compartments of skeletal muscle fibres.

## Acknowledgements

The imaging by transmission electron microscopy was performed at the Core Facility for Integrated Microscopy, Faculty of Health and Medical Science, University of Copenhagen.

## Additional information

## Data availability statement

Raw images, annotations, and model weights are available here: https://doi.org/10.5281/zenodo.18390286. The code for traning, testing, and a graphical user interface can be found here: https://github.com/JoachimNielsenSDU/Glycogen-segmenter.

## Author contributions

AAH, JME, KS, DW, JJ, and JN contributed to the conception and design of the experiments. All authors contributed to the data collection, analyses and/or data interpretation. AAH, JME, and JN drafted the manuscript, while all authors edited and/or revised the manuscript. The final version of the manuscript was approved by all authors, and all agree to be accountable for all aspects of the work in ensuring that questions related to the accuracy or integrity of any part of the work are appropriately investigated and resolved. All persons stated as authors qualify for authorship, and all those who qualify for authorship are listed.

